# Genomic evidence of globally widespread admixture from polar bears into brown bears during the last ice age

**DOI:** 10.1101/154773

**Authors:** James A. Cahill, Peter D. Heintzman, Kelley Harris, Matthew Teasdale, Joshua Kapp, André E Rodrigues Soares, Ian Stirling, Daniel Bradley, Ceiridwen J. Edwards, Aliaksandr A. Kisleika, Alexander V. Malev, Nigel Monaghan, Richard E. Green, Beth Shapiro

**Affiliations:** Department of Ecology and Evolutionary Biology, University of California Santa Cruz, Santa Cruz, CA 95064, USA; Tromsø University Museum, UiT - The Arctic University of Norway, 9037 Tromsø, Norway.; Department of Genetics, Stanford University, Stanford, CA 94305; Smurfit Institute of Genetics, Trinity College, Dublin, Ireland.; Wildlife Research Division, Department of Environment, c/o Department of Biological Sciences, University of Alberta, Edmonton, AB, T6G 2E9; Department of Biological Sciences, University of Alberta, Edmonton, AB, TGH 2E9; Department of Biological Sciences, School of Applied Sciences, University of Huddersfield, Queensgate, Huddersfield, HD1 3DH, UK; Director of Kurils Nature Reserve, Russia, Yuzhno-Kurilsk, 694500 Zarechnaya str., 5; Russia, Kazan, 420059, Khadi Taktash str. 112, Deputy director of Kazan Zoobotanical garden; National Museum of Ireland – Natural History, Merrion Street, Dublin D22 F627, Ireland; Department of Biomolecular Engineering, University of California Santa Cruz, Santa Cruz, CA 95064, USA; UCSC Genomics Institute, University of California Santa Cruz, Santa Cruz, CA 95064, USA

## Abstract

Recent genomic analyses have provided substantial evidence for past periods of gene flow from polar bears (*Ursus maritimus*) into Alaskan brown bears (*Ursus arctos*), with some analyses suggesting a link between climate change and genomic introgression. However, because it has only been possible to sample bears from the present day, the timing, frequency, and evolutionary significance of this admixture remains unknown. Here, we analyze genomic DNA from three additional and geographically distinct brown bear populations, including two that lived temporally close to the peak of the last ice age. We find evidence of admixture in all three populations, suggesting that admixture between these species has been common in their recent evolutionary history. In addition, analyses of ten fossil bears from the now-extinct Irish population indicate that admixture peaked during the last ice age, when brown bear and polar bear ranges overlapped. Following this peak, the proportion of polar bear ancestry in Irish brown bears declined rapidly until their extinction. Our results support a model in which ice age climate change created geographically widespread conditions conducive to admixture between polar bears and brown bears, as is again occurring today. We postulate that this model will be informative for many admixing species pairs impacted by climate change. Our results highlight the power of paleogenomes to reveal patterns of evolutionary change that are otherwise masked with only contemporary data.

## Introduction

Post-divergence gene flow between species is increasingly understood to have been common in evolutionary history (Green et al. 2010; Dasmahapatra et al. 2012; Poelstra et al. 2014; Lamichhaney et al. 2015). Also known as admixture, this process most commonly occurs when two formerly geographically isolated species overlap in range and are reproductively compatible. Genomic analyses across hybrid zones have revealed considerable variation among species pairs in both the spatial patterns and evolutionary consequences of admixture (Good et al. 2008; Dasmahapatra et al. 2012; Poelstra et al. 2014). In some cases, genomic incompatibilities lead to hybrid phenotypes that are less fit than either parent species (Good et al. 2008). In other cases, new combinations of alleles may provide local adaptive advantages (Garroway et al. 2010). Hybridization may, therefore, be an important source of evolutionary novelty, for example during periods of rapid climate change, when shifting habitats may form communities comprising previously isolated populations and species (Graham et al. 1996; Parmesan and Yohe 2003; Hoffmann and Sgrò 2011).

Polar bears and brown bears diverged less than 500 thousand years ago (Hailer et al. 2012; Cahill et al. 2013; Liu et al. 2014) but differ morphologically, physiologically, and behaviorally (Stirling 2011). In recent years, whole genome sequencing has revealed that all North American brown bears have between 3% and 8% of their genome derived from polar bear ancestry (Cahill et al. 2013; Liu et al. 2014; Cahill et al. 2015). Polar bear ancestry is greatest among North American brown bears in Southeast Alaska’s ABC (Admiralty, Baranof and Chichagof) Islands (Liu et al. 2014; Cahill et al. 2015). Previously, we proposed a population conversion model of polar/brown bear admixture (Cahill et al. 2013), in which a warming climate at the end of the last ice age allowed brown bears to disperse into what had previously been polar bear range (today’s ABC Islands), resulting in hybridization and the formation of a hybrid population (Cahill et al. 2013; Cahill et al. 2015). Even after the climate stabilized during the Holocene, the ABC Islands continued to receive immigrants from the much larger and less admixed mainland population, which gradually reduced the polar bear contribution in the population to the 6-8% observed in ABC islands brown bears today (Cahill et al. 2015).

An alternative hypothesis has also been proposed, however. Liu *et* al (Liu et al. 2014) suggested that the admixture event occurred prior to the last ice age, and that the proportion of polar bear ancestry in ABC Islands bears has been relatively stable since that time. These models make very different predictions about the impact of admixture on populations’ genetic diversity, with the population conversion model suggesting a much stronger impact than the alternative model, and about the subsequent fate of that diversity, with the population conversion model supporting a gradual loss of diversity over time. Both models are consistent with some features of the nuclear genomic data from present-day individuals, and so the absence of a direct measurement of polar bear ancestry in the past has prevented resolution of this question.

Here, we use a paleogenomic approach to directly explore the role of climate change in facilitating admixture between brown bears (*Ursus arctos*) and polar bears (*U. maritimus*) (Fig. 1). Focusing on a now-extinct population of brown bears from Ireland, we isolated genomic DNA from ten cave-preserved bones that were morphologically and isotopically identified as brown bears (Edwards et al. 2011) and that range in age from 37.5-3.9 thousand calibrated years before present (cal. ka BP). This interval spans the local peak of the last ice age *ca.* 24.7 cal. ka BP (Peters et al. 2015), when polar bears’ distribution would have been most proximate to present-day Ireland. Previously, mitochondrial DNA showed that some Irish brown bear fossils have polar bear-like mitochondrial haplotypes, which is consistent with admixture having occurred between polar bears and brown bears in Ireland (Edwards et al. 2011).

**Fig. 1.**
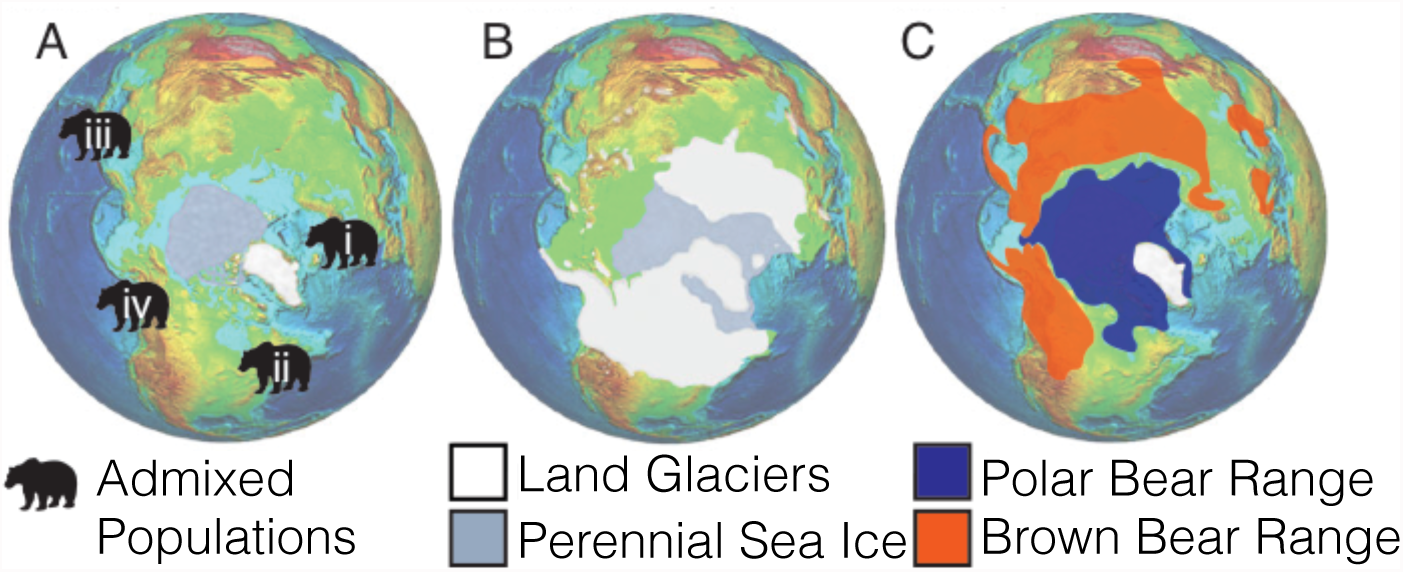
(A) Geographic locations of brown bear populations identified here and in previous analyses (Cahill et al. 2013) as having some component of polar bear ancestry: (i) present-day Ireland; (ii) Chaplain Sea, Quebec, Canada; (iii) Kunashir Island, Russia; (iv) Admiralty, Baranof and Chichagof (ABC) Islands, Alaska, USA. Panel A shows the present day distribution of glaciers and sea ice. Details of samples used here are provided in supplementary table S3. Each of these admixed populations is located near the extent of sea and/or glacial ice at the last glacial maximum, *ca.* 24 ka BP (Peters et al. 2015), which is depicted in panel B, but far from the present-day range of polar bears (Schliebe et al. 2008), as shown in panel C. Base image from (http://earthobservatory.nasa.gov/Features/BorealMigration/boreal_migration2.php).

To explore the geographic extent of potential admixture between brown bears and polar bears, we also extracted and analyzed DNA from an 11.3 cal. ka BP (Harington et al. 2014) brown bear bone from the coast of the Champlain Sea in Quebec, Canada, and from two bears from the present-day population of Kunashir Island, in eastern Russia (Fig. 1). Both populations were located near perennial sea ice during the Last Glacial maximum (LGM) (Seki et al. 2004; Harington 2008), suggesting they may have been regions where polar bear and brown bear ranges overlapped. Interestingly, some Kunashir Island brown bears have white coats (Sato et al. 2011) and, while the source of this color variation is unknown, intermediate coat color is typical of polar/brown bear hybrids (Preuß et al. 2009). These samples, together with the Irish brown bears and the ABC islands brown bears, provide one population each from the east and west coasts of the Atlantic and Pacific, allowing us to test whether admixture between polar bears and brown bears was constrained to a small number of islands or was widespread throughout the Northern Temperate zone.

## Results

We used the *D* and 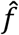 statistics (Green et al. 2010; Durand et al. 2011), to estimate the amount of polar bear ancestry within each Irish bear genome (fig, 2, supplementary Fig. S1,S2, supplementary table S1, S2). To assess statistical significance, we used the weighted block jackknife with 5Mb blocks (Kunsch 1989). Z-scores are calculated by dividing the D or 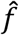 value by the weighted block jackknife standard error. Z-scores greater than 3 are considered significant evidence of admixture. We found all 10 Irish brown bears to have significant polar bear ancestry ranging from at least 3% to at most 21.5% of their genomes (Fig. 2, supplementary Fig. S1; supplementary table S1, S2). Strikingly, the Irish brown bears with the largest proportion of polar bear ancestry lived temporally closest to the peak of the last ice age, with the most admixed bear, 21.5% polar bear ancestry (Z=11.7), dating to *ca.* 13 cal. ka BP. Observed polar bear ancestry in Irish brown bears generally declined between 13 cal. ka BP and 4 cal. ka BP (Fig. 2).

**Fig. 2.**
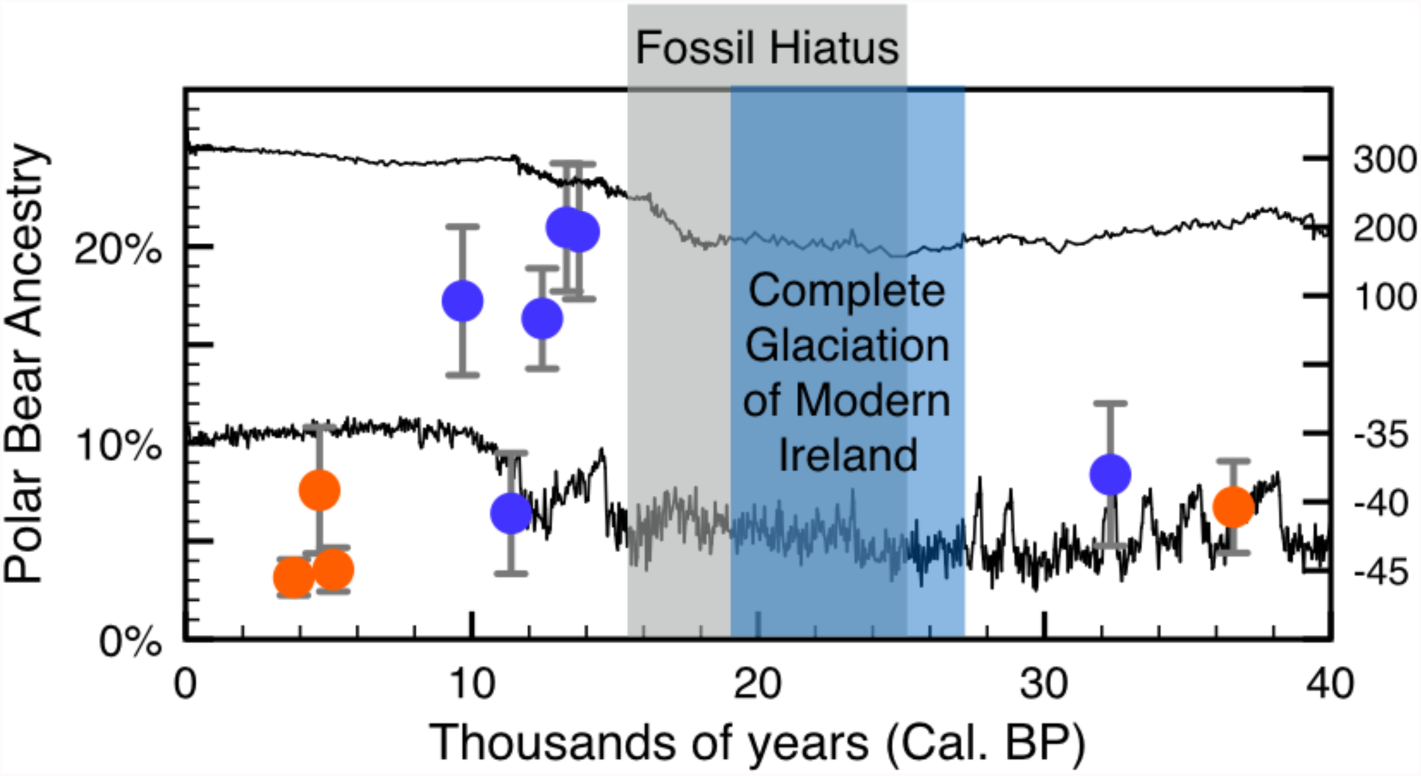
The percentage of each Irish brown bear genome derived from polar bear ancestry, estimated using 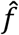 and plotted against its calibrated age (Methods). Error bars show 95% confidence intervals estimated by weighted block jackknife (1.96 standard errors). Mitochondrial haplotype (Edwards et al. 2011) is indicated by color: polar bear like, clade 2 (blue) and brown bear like, clade 1 (orange). To show the correspondence between polar bear ancestry and climate, we show two climate proxies: δO^18^ from NGRIP and CO_2_ from Vostok; in both cases, values closer to the top of the Fig. indicate warmer temperatures. Glacial reconstructions indicate that all of modern Ireland was glaciated during the local peak of the last ice age from 27-19 ka (Clark et al. 2012), although radiocarbon dates indicate that some areas in the far south-east may have been ice free as late as 25 ka BP (Woodman et al. 1997). A general hiatus in the vertebrate fossil record is known in Ireland from the glacial peak until 15 ka BP (Woodman et al. 1997; Stuart et al. 2004). Brown bears occur in the Irish fossil record both before and after the glacial peak, but are absent during from 32-14 ka BP (Woodman et al. 1997; Edwards et al. 2011). The most recent pre-glacial and most ancient post-glacial brown bear bones from the Irish fossil record are analyzed as part of this study.

The two other previously unstudied populations were also found to exhibit polar bear ancestry. The brown bears from Kunashir Island in Eastern Russia have 4.0% (Z=6.1) and 12.7% (Z=16.9) polar bear ancestry (supplementary Fig. S1; supplementary table S1, S2), strongly supporting past admixture with polar bears. Likewise the 11.3 cal. ka BP (Harington et al. 2014) brown bear bone recovered from Champlain Sea deposits in Quebec, Canada (Fig. 1) has at least 8.5% (Z=5.7) polar bear ancestry (supplementary Fig. S1; supplementary table S1, S2).

The D and 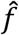 statistics test for the presence of admixture but do not explicitly test whether that is gene flow from polar bears into brown bears or the reverse (Green et al. 2010; Durand et al. 2011). In all three candidate hybrid populations (Ireland, Kunashir and Quebec), we tested the direction of gene flow by scanning the genome for regions where putative hybrids were excessively similar to polar bears and divergent from other brown bears, which would indicate the introgression of polar bear alleles into the hybrid brown bears (Supplementary Material online)(Green et al. 2010). Consistent with results in previously studied admixed bear populations (Cahill et al. 2013; Liu et al. 2014) we found the candidate hybrids to be enriched for genomic regions of low polar bear divergence and high brown bear divergence relative to a Fennoscandian brown bear with no detectable polar bear ancestry (supplementary Fig. S3). This provides additional support for polar bear introgression into the Ireland, Kunashir and Quebec brown bear populations.

The marked decline in Irish brown bears’ polar bear ancestry from 14,000 to 3,000 years ago could be driven by: demographic processes, such as the immigration of unadmixed brown bears decreasing the fraction of polar bear ancestry in the population; selection against polar bear alleles; or some combination of the two. The very small effective population size of polar bears (Miller et al. 2012) may have led to an overall higher genetic load compared to brown bears, which would exert selection against polar bear ancestry. To test whether selection against maladaptive polar bear alleles contributed to the post-glacial decline in polar bear ancestry, we performed simulations of the impact of selection against introgressed loci due to polar bears’ greater genetic load under a recently proposed population history model (Liu et al. 2014) (see Methods). Our simulations show that polar bears are expected to have only a slightly higher genetic load than brown bears, resulting in a median decrease in fitness of 4% (supplementary Fig. S4). This resulted in a simulated decline in polar bear ancestry, from a starting value of 25%, to 22.4% before stabilizing (supplementary Fig. S5). The post-LGM reduction in polar bear ancestry is, therefore, not primarily explained by genetic load or selection against polar bear alleles.

The samples from Ireland were preserved in temperate caves and exhibit degradation and low endogenous DNA content typical of such samples (supplementary table S3, S4). A pilot study showed that the Irish brown bear samples have the best preserved nuclear DNA of any Irish brown bear bones in the National Museum of Ireland Collection (supplementary online materials), however, achieving 1x coverage for any, let alone all, of the samples is not feasible given the DNA preservation. Although, the coverage is <1x it is sufficiently high that all of the Irish brown bears were found to have significant polar bear ancestry (Z>3).

To test whether the methods in this study were robust at the coverage that was attainable from the samples we simulated low coverage samples randomly sampled reads from a multi-fold coverage modern brown bear known to have 8.56% autosomal polar bear ancestry (Cahill et al. 2015) (supplementary table S5). We found lower coverage samples are more susceptible to ascertainment bias from reference genome selection, resulting in overestimation of polar bear ancestry, than higher coverage samples but this can be mitigated by appropriate read mapping parameter selection (supplementary Fig. S6). Ten simulations of 1 million randomly sampled mapped reads (fewer reads than 4 of the Irish bears and both Kunashir bears), produced estimates from 8.50% to 9.70% with a mean of 9.04% (supplementary Fig. S6, supplementary table S5). That this previously undocumented bias in D and f statistic estimation exists should serve as a caution to future studies, however, 0.48% is not comparable to the magnitude of the admixture observed in this study. Further, the narrow range of estimates in the test, 1.20% between the most extreme results for 1 million mapped reads (supplementary table S5), show that the coverage used in this study can accurately and reliably estimate introgressed ancestry.

## Discussion

The very high polar bear ancestry observed immediately after the local peak of the last ice age was followed by a decline in polar bear ancestry. This closely corresponds to the population conversion model previously proposed for admixture between polar bears and brown bears on southeastern Alaska’s ABC Islands (Cahill et al. 2013). That the population conversion model is operative in Ireland suggests that it was also operative in the ABC Islands. We further suggest that the population conversion model predicted from ABC Islands genomic diversity and directly observed in the Irish population may be a general pattern for polar/brown bear admixture.

We hypothesize that a combination of local changes in habitat availability, lower sea levels, and species-specific natural histories facilitated the observed admixture between brown and polar bears. The paleoecological and fossil records of Ireland suggest that all or most of Ireland was glaciated throughout much of the last ice age, leaving little to no habitat for brown bears (Clark et al. 2012; Ó Cofaigh et al. 2012; Edwards et al. 2014). At the same time, major tidewater glaciers on the western shelf and down the Irish Sea basin and offshore iceberg scouring of the sea floor suggest the possibility of productive sea ice habitat for polar bears along the Irish coast (Edwards et al. 2011; Clark et al. 2012; Ó Cofaigh et al. 2012; Peters et al. 2015). As resident brown bear populations declined during the approach of the LGM, this proximity in range probably led to admixture, as it can in present day populations of brown bears and polar bears whose ranges overlap (Stirling 2011). After the LGM, brown bears likely recolonized Ireland from mainland Europe or Great Britain (Edwards et al. 2014). These colonizing bears would have encountered and hybridized with resident polar bears or hybrid bears. Such encounters probably decreased in frequency as the ice receded. Continuing dispersal from the mainland of non-admixed brown bears would reduce the proportion of polar bear ancestry in the Irish brown bear population, leading to the pattern observed in the Irish brown bear genomes (Fig. 2).

The observation that the Kunashir and Quebec populations also have polar bear ancestry provides further support for the conclusion that the ABC islands and Ireland are part of a broader pattern of admixture and not isolated idiosyncratic events. Kunashir Island is the first Asian brown bear population shown to have polar bear ancestry. This population may include individuals with greater amounts of polar bear ancestry than the ABC Islands brown bears, as the Kunashir 2 individual’s 12% polar bear ancestry exceeds the 8% that was the most polar bear ancestry previously observed in an extant brown bear (Cahill et al. 2015). This may reflect a different demographic or selective regime in the Kunashir Islands than in the ABC Islands, which could be explored in future research. However, we suggest some caution in interpreting this result, because the Kunashir 2 value varies more than others according to the choice of bioinformatic approach (supplementary Fig. S1, S2). Nonetheless, all of our analyses support both Kunashir bears as having polar bear ancestry (supplementary table S1,S2).

Admixture between brown bears and polar bears has also been observed in the present-day Canadian Arctic (Doupé et al. 2007; Pongracz et al. 2017) and has been attributed to climate-induced overlap between the two species (Kelly et al. 2010). Together, these data reveal the ongoing and dynamic nature of gene flow between brown bears and polar bears, and the important role that climate change and consequent habitat redistribution plays in facilitating admixture. Intriguingly, the evolutionary consequences of this admixture appear to be mediated by ecological and behavioral differences between the two species, which maintain polar bears as a genetically distinct lineage that lacks any detectable of brown bear introgression (Cahill et al. 2015; Peacock et al. 2015). These results highlight the complicated nature of speciation, and suggest that *Ursus*, which includes brown bears and polar bears, may be a useful genus in which to explore the formation of incompatibilities between diverging lineages.

## Conclusion

Admixture between polar bears and brown bears is geographically widespread, and associated with fluctuations in climate surrounding the last ice age and the present warming period. In Ireland, the proportion of polar bear ancestry in resident brown bears peaked after the last ice age and then declined until the population’s extinction ~4 cal. ka BP (Fig. 2). This pattern is consistent with the population conversion model of admixture previously suggested to explain the extant admixed population on Alaska’s ABC Islands (Cahill et al. 2013), and highlights the power of paleogenomics to test demographic and evolutionary hypotheses.

Correlation between recent climate change and admixture has been observed recently for several related species pairs, including trout (Muhlfeld et al. 2014), flying squirrels (Garroway et al. 2010), *Pachycladon* grasses (Becker et al. 2013), and damselflies (Sánchez-Guillén et al. 2014). Although the long-term evolutionary consequences to these species pairs are not yet known, preliminary evidence suggests a wide range of possible outcomes, from extinction via genetic replacement (Muhlfeld et al. 2014) to the creation of hybrid phenotypes with higher fitness in the new habitat relative to the parental lineages (Becker et al. 2013). While it is tempting to consider these as localized examples, and therefore unlikely to have widespread evolutionary consequences, introgressed DNA will in many instances spread to non-admixed populations as individuals disperse. For example, introgressed polar bear DNA has been observed in mainland Alaskan brown bears, probably due to post-glacial dispersal from the ABC Islands (Liu et al. 2014; Cahill et al. 2015). Thus admixture resulting from climate-related habitat redistribution is likely to have long-term and widespread evolutionary consequences, and may be an important mechanism for generating and maintaining diversity.

## Methods

### DNA extraction, library preparation and sequencing

All pre-amplification laboratory work on the ancient specimens was conducted in a dedicated clean lab facility at the UC Santa Cruz Paleogenomics Lab, following standard procedures for working with degraded DNA (Fulton 2012). We tested multiple extraction methods to optimize DNA recovery (Supplementary Material online). After DNA extraction, we converted the extracts into double-stranded, indexed sequencing libraries following (Meyer and Kircher 2010), as modified by (Heintzman et al. 2015). We then pooled the libraries and sequenced them on the Illumina MiSeq and HiSeq 2500 platforms (Supplementary Material online).

### Mapping and reference bias correction

To identify optimal mapping parameters for this data we compared a range of parameters to an optimal read alignment (Supplementary Material online, supplementary table S6). This led us to select bowtie2 v2.1.0 (Langmead and Salzberg 2012), with the local alignment approach (-local flag), allowing a single mismatch allowed in the mapping seed (-N 1 flag) and a maximum mismatch penalty of 4 (-mp 4 flag) for use with the Ireland, Quebec and Kunashir samples. We excluded read mappings with map quality scores <30 and removed duplicate reads with samtools v0.1.19 (Li et al. 2009).

To describe and mitigate the impact of ascertainment bias from the reference genome we mapped reads to the polar bear reference genome (Liu et al. 2014) and a consensus sequence of an unadmixed Swedish brown bear (SJS01)(Liu et al. 2014). As expected each genome produced a slightly different set of read mappings (supplementary table S4). Mappings to the polar bear reference results in greater inferred polar bear introgression the mapping to brown bear consensus sequence (supplementary Fig. S1) suggesting that both methods are somewhat biased toward their respective species. To minimize bias and capture as much of the admixed bears’ genomes as possible we combined the mappings to the polar bear and brown bear references and for each read retained the mapping coordinates with the highest map quality score using an in house script (to be released on GitHub) (Supplementary Material online). This two reference approach recovered more reads than either single reference approach indicating that both references must have contributed unique mappings (supplementary table S4) and produced intermediate estimates of polar bear ancestry (supplementary Fig. S1, supplementary tables S1,S2) which we consider to be minimally biased.

### Estimating the proportion of polar bear ancestry

We used the D-statistic (also known as the ABBA/BABA test) to test for the possibility of admixture between polar bears and the Ireland, Quebec and Kunashir Island brown bears (Green et al. 2010; Durand et al. 2011). For our comparisons we considered each of 30 polar bears (Miller et al. 2012; Cahill et al. 2013) and 4 Fennoscandian brown bears (Liu et al. 2014; Cahill et al. 2015), all of which had been previously shown to lack detectable introgressed ancestry (Liu et al. 2014; Cahill et al. 2015). To identify the ancestral state we used an American black bear (*Ursus americanus*) and a giant panda (*Ailuropoda melanoleuca*) as outgroups. The two outgroups produced similar admixture estimates (supplementary Fig. S2) so we used the more closely related American black bear for all remaining analyses. To quantify the amount of admixture we used the 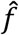 statistic which compares the observed derived allele sharing between a hybrid and a polar bear with the amount of derived allele sharing expected between two polar bears, conceptually a 100% introgressed hybrid (Green et al. 2010; Durand et al. 2011)( Supplementary Material online), For both D and 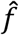 we tested the impact of different reference genomes, and the inclusion or exclusion of transition sites which are susceptible to ancient DNA damage induced bias (supplementary Fig. S1, Supplementary Material online) (Hofreiter 2001). We test for significant evidence of admixture with the weighted block jackknife (Kunsch 1989) with 5 Mb non-overlapping blocks, we consider Z-scores>3 to be significant. We tested the direction of gene flow with the method of (Green et al. 2010)(supplementary Fig. S3, Supplementary Material online), and test for unexpected biases resulting from less than 1x coverage by randomly sampling reads from a high coverage modern brown bear with known polar bear ancestry (Supplementary Material online).

### Influence of selection against polar bear alleles in hybrids

To test whether the accumulation of weakly deleterious alleles in polar bears could be responsible for the decline in polar bear ancestry observed in the Irish brown bear population (Fig. 2), we used the forward-simulation approach of (Harris and Nielsen 2016), and the simulator SLiM (Messer 2013). For this analysis, we drew model parameters from a recent inference of polar bear demographic history (Liu et al. 2014)(Supplementary Materials). To model the admixture occurring in Ireland we simulated a single 25% pulse of polar bear introgression into a brown bear population 15,000 years ago. We measured the expected difference in fitness between polar bears and brown bears under the model at the time of the simulated admixture (supplementary Fig. S4) and the change in polar bear ancestry in the admixed population resulting from greater genetic load (supplementary Fig. S5).

## Data Accessibility

Raw sequence reads will be released on NCBI at the time of publication. Programs not previously published will be released on GitHub.

## References

Becker M, Gruenheit N, Steel M, Voelckel C, Deusch O, Heenan PB, McLenachan PA, Kardailsky O, Leigh JW, Lockhart PJ. 2013. Hybridization may facilitate in situ survival of endemic species through periods of climate change. Nat. Clim. Chang. [Internet] 3:1039–1043. Available from: http://www.nature.com/doifinder/10.1038/nclimate2027

Cahill JA, Green RE, Fulton TL, Stiller M, Jay F, Ovsyanikov N, Salamzade R, St John J, Stirling I, Slatkin M, et al. 2013. Genomic evidence for island population conversion resolves conflicting theories of polar bear evolution. Nachman MW, editor. PLoS Genet. [Internet] 9:e1003345. Available from: http://dx.plos.org/10.1371/journal.pgen.1003345

Cahill JA, Stirling I, Kistler L, Salamzade R, Ersmark E, Fulton TL, Stiller M, Green RE, Shapiro B. 2015. Genomic evidence of geographically widespread effect of gene flow from polar bears into brown bears. Mol. Ecol. [Internet] 24:1205–1217. Available from: http://www.ncbi.nlm.nih.gov/pubmed/25490862

Clark CD, Hughes ALC, Greenwood SL, Jordan C, Sejrup HP. 2012. Pattern and timing of retreat of the last British-Irish Ice Sheet. Quat. Sci. Rev. [Internet] 44:112–146. Available from: http://linkinghub.elsevier.com/retrieve/pii/S0277379110002817

Dasmahapatra KK, Walters JR, Briscoe AD, Davey JW, Whibley A, Nadeau NJ, Zimin A V., Hughes DST, Ferguson LC, Martin SH, et al. 2012. Butterfly genome reveals promiscuous exchange of mimicry adaptations among species. Nature [Internet] 487:94–98. Available from: http://www.nature.com/doifinder/10.1038/nature11041

Doupé JP, England JH, Furze M, Paetkau D. 2007. Most Northerly Observation of a Grizzly Bear (Ursus arctos) in Canada: Photographic and DNA Evidence from Melville Island, Northwest Territories. Arctic [Internet] 60:271–276. Available from: http://www.jstor.org/stable/40512895?seq=1#page_scan_tab_contents

Durand EY, Patterson N, Reich D, Slatkin M. 2011. Testing for ancient admixture between closely related populations. Mol. Biol. Evol. [Internet] 28:2239–2252. Available from: http://mbe.oxfordjournals.org/content/28/8/2239.full

Edwards CJ, Ho SYW, Barnett R, Coxon P, Bradley DG, Lord TC, O’Connor T. 2014. Continuity of brown bear maternal lineages in northern England through the Last-glacial period. Quat. Sci. Rev. [Internet] 96:131–139. Available from: http://www.sciencedirect.com/science/article/pii/S0277379113004095

Edwards CJ, Suchard MA, Lemey P, Welch JJ, Barnes I, Fulton TL, Barnett R, O’Connell TC, Coxon P, Monaghan N, et al. 2011. Ancient hybridization and an Irish origin for the modern polar bear matriline. Curr. Biol. [Internet] 21:1251–1258. Available from: http://www.ncbi.nlm.nih.gov/pubmed/21737280

Fulton TL. 2012. Setting up an ancient DNA laboratory. In: Shapiro B, Hofreiter M, editors. Ancient DNA, methods and protocols. Springer protocols. p. 1–11.

Garroway CJ, Bowman J, Cascaden TJ, Holloway GL, Mahan CG, Malcolm JR, Steele MA, Turner G, Wilson PJ. 2010. Climate change induced hybridization in flying squirrels. Glob. Chang. Biol. [Internet] 16:113–121. Available from: http://doi.wiley.com/10.1111/j.1365-2486.2009.01948.x

Good JM, Dean MD, Nachman MW. 2008. A complex genetic basis to X-linked hybrid male sterility between two species of house mice. Genetics [Internet] 179:2213–2228. Available from: http://www.genetics.org/content/179/4/2213.short

Graham RW, Lundelius EL, Graham MA, Schroeder EK, Toomey RS, Anderson E, Barnosky AD, Burns JA, Churcher CS, Grayson DK, et al. 1996. Spatial Response of Mammals to Late Quaternary Environmental Fluctuations. Science (80-.). 272:1601–1606.

Green RE, Krause J, Briggs AW, Maricic T, Stenzel U, Kircher M, Patterson N, Li H, Zhai W, Fritz MH-Y, et al. 2010. A draft sequence of the Neandertal genome. Science [Internet] 328:710–722. Available from: http://www.sciencemag.org/content/328/5979/710.abstract

Hailer F, Kutschera VE, Hallström BM, Klassert D, Fain SR, Leonard JA, Arnason U, Janke A. 2012. Nuclear genomic sequences reveal that polar bears are an old and distinct bear lineage. Science [Internet] 336:344–347. Available from: http://www.ncbi.nlm.nih.gov/pubmed/22517859

Harington CR. 2008. The evolution of Arctic marine mammals. Ecol. Appl. 18:S23–S40.

Harington CR, Cournoyer M, Chartier M, Fulton TL, Shapiro B. 2014. Brown bear (Ursus arctos) (9880 ± 35bp) from late-glacial Champlain Sea deposits at Saint-Nicolas, Quebec, Canada, and the dispersal history of brown bears. Can. J. Earth Sci. 535:527–535.

Harris K, Nielsen R. 2016. The Genetic Cost of Neanderthal Introgression. Genetics [Internet]: genetics. 116.186890-. Available from: http://www.genetics.org/content/early/2016/03/31/genetics.116.186890

Heintzman PD, Zazula GD, Cahill JA, Reyes A V., MacPhee RDE, Shapiro B. 2015. Genomic Data from Extinct North American Camelops Revise Camel Evolutionary History. Mol. Biol. Evol. [Internet] 32:2433–2440. Available from: http://mbe.oxfordjournals.org/lookup/doi/10.1093/molbev/msv128

Hoffmann AA, Sgró CM. 2011. Climate change and evolutionary adaptation. Nature [Internet] 470:479–485. Available from: http://www.nature.com/nature/journal/v470/n7335/full/nature09670.html#ref70

Hofreiter M. 2001. DNA sequences from multiple amplifications reveal artifacts induced by cytosine deamination in ancient DNA. Nucleic Acids Res. [Internet] 29:4793–4799. Available from: http://nar.oxfordjournals.org/content/29/23/4793

Kelly BP, Whiteley A, Tallmon D. 2010. The Arctic melting pot. Nature 468:891.

Kunsch HR. 1989. The Jackknife and the Bootstrap for General Stationary Observations. Ann. Stat. [Internet] 17:1217–1241. Available from: http://projecteuclid.org/euclid.aos/1176347265

Lamichhaney S, Berglund J, Almén MS, Maqbool K, Grabherr M, Martinez-Barrio A, Promerová M, Rubin C-J, Wang C, Zamani N, et al. 2015. Evolution of Darwin’s finches and their beaks revealed by genome sequencing. Nature [Internet] 518:371–375. Available from: http://dx.doi.org/10.1038/nature14181

Langmead B, Salzberg SL. 2012. Fast gapped-read alignment with Bowtie 2. Nat. Methods [Internet] 9:357–359. Available from: http://dx.doi.org/10.1038/nmeth.1923

Li H, Handsaker B, Wysoker A, Fennell T, Ruan J, Homer N, Marth G, Abecasis G, Durbin R. 2009. The Sequence Alignment/Map format and SAMtools. Bioinformatics [Internet] 25:2078–2079. Available from: http://bioinformatics.oxfordjournals.org/content/25/16/2078.short

Liu S, Lorenzen ED, Fumagalli M, Li B, Harris K, Xiong Z, Zhou L, Korneliussen TS, Somel M, Babbitt C, et al. 2014. Population Genomics Reveal Recent Speciation and Rapid Evolutionary Adaptation in Polar Bears. Cell [Internet] 157:785–794. Available from: http://www.cell.com/article/S0092867414004887/fulltext

Messer PW. 2013. SLiM: Simulating evolution with selection and linkage. Genetics 194:1037–1039.

Meyer M, Kircher M. 2010. Illumina sequencing library preparation for highly multiplexed target capture and sequencing. Cold Spring Harb. Protoc. [Internet] 2010:pdb.prot5448. Available from: http://www.ncbi.nlm.nih.gov/pubmed/20516186

Miller W, Schuster SC, Welch AJ, Ratan A, Bedoya-Reina OC, Zhao F, Kim HL, Burhans RC, Drautz DI, Wittekindt NE, et al. 2012. Polar and brown bear genomes reveal ancient admixture and demographic footprints of past climate change. Proc. Natl. Acad. Sci. U. S. A. [Internet] 109:E2382-90. Available from: http://www.pubmedcentral.nih.gov/articlerender.fcgi?artid=3437856&tool=pmcentrz&rendertype=abstract

Muhlfeld CC, Kovach RP, Jones LA, Al-Chokhachy R, Boyer MC, Leary RF, Lowe WH, Luikart G, Allendorf FW. 2014. Invasive hybridization in a threatened species is accelerated by climate change. Nat. Clim. Chang. [Internet] 4:620–624. Available from: http://dx.doi.org/10.1038/nclimate2252

Ó Cofaigh C, Telfer MW, Bailey RM, Evans DJA. 2012. Late Pleistocene chronostratigraphy and ice sheet limits, southern Ireland. Quat. Sci. Rev. [Internet] 44:160–179. Available from: http://www.sciencedirect.com/science/article/pii/S0277379110000132

Parmesan C, Yohe G. 2003. A globally coherent fingerprint of climate change impacts across natural systems. Nature [Internet] 421:37–42. Available from: http://dx.doi.org/10.1038/nature01286

Peacock E, Sonsthagen SA, Obbard ME, Boltunov A, Regehr E V, Ovsyanikov N, Aars J, Atkinson SN, Sage GK, Hope AG, et al. 2015. Implications of the circumpolar genetic structure of polar bears for their conservation in a rapidly warming arctic. PLoS One [Internet] 10:e112021. Available from: http://journals.plos.org/plosone/article?id=10.1371/journal.pone.0112021#pone-0112021-g003

Peters JL, Benetti S, Dunlop P, Ó Cofaigh C. 2015. Maximum extent and dynamic behaviour of the last British–Irish Ice Sheet west of Ireland. Quat. Sci. Rev. [Internet] 128:48–68. Available from: http://linkinghub.elsevier.com/retrieve/pii/S027737911530113X

Poelstra JW, Vijay N, Bossu CM, Lantz H, Ryll B, Müller I, Baglione V, Unneberg P, Wikelski M, Grabherr MG, et al. 2014. The genomic landscape underlying phenotypic integrity in the face of gene flow in crows. Science [Internet] 344:1410–1414. Available from: http://www.sciencemag.org/content/344/6190/1410.short

Pongracz JD, Paetkau D, Branigan M, Richardson E. 2017. Recent Hybridization between a Polar Bear and Grizzly Bears in the Canadian Arctic. Arctic 70.

Preuß A, Gansloßer U, Purschke G, Magiera U. 2009. Bear-hybrids: behaviour and phenotype. Der Zool. Garten [Internet] 78:204–220. Available from: http://www.sciencedirect.com/science/article/pii/S0044516909000276

Sánchez-Guillén RA, Muñoz J, Hafernik J, Tierney M, Rodriguez-Tapia G, Córdoba-Aguilar A. 2014. Hybridization rate and climate change: are endangered species at risk? J. Insect Conserv. [Internet] 18:295–305. Available from: http://link.springer.com/10.1007/s10841-014-9637-5

Sato Y, Nakamura H, Ishifune Y, Ohtaishi N. 2011. The white-colored brown bears of the Southern Kurils. Ursus 22:84–90.

Schliebe S, Wiig Ø, Derocher A, Lunn N, (IUCN SSC Polar Bear Specialist Group). 2008. Ursus maritimus (Polar Bear). IUCN Red List Threat. Species. Version 2015.2. [Internet]. Available from: http://www.iucnredlist.org/details/22823/0

Seki O, Ikehara M, Kawamura K, Nakatsuka T, Ohnishi K, Wakatsuchi M, Narita H, Sakamoto T. 2004. Reconstruction of paleoproductivity in the Sea of Okhotsk over the last 30 kyr. Paleoceanography Available from: http://doi.wiley.com/10.1029/2002PA000808

Stirling I. 2011. Polar Bears: The Natural History of a Threatened Species. Brighton, MA: Fitzhenry and Whiteside

Stuart AJ, Kosintsev PA, Higham TFG, Lister AM. 2004. Pleistocene to Holocene extinction dynamics in giant deer and woolly mammoth. Nature [Internet] 431:684–689. Available from: http://dx.doi.org/10.1038/nature02890

Woodman P, McCarthy M, Monaghan N. 1997. The Irish Quaternary Fauna Project. Quat. Sci. Rev. 16:129–159.

